# Cyclophilin D induces necrotic core formation by promoting mitochondria-mediated macrophage apoptosis in advanced atherosclerotic lesions

**DOI:** 10.1101/2023.09.05.556288

**Authors:** Jun-ichiro Koga, Ryuta Umezu, Tomohiro Shirouzu, Yuki Kondo, Nasanbadrakh Orkhonselenge, Hiromichi Ueno, Shunsuke Katsuki, Tetsuya Matoba, Yosuke Nishimura, Masaharu Kataoka

## Abstract

**Background:** In advanced atherosclerotic lesions, apoptotic cell death of plaque macrophages results in necrotic core formation and plaque vulnerability. Cyclophilin D (CypD) is a mitochondria-specific cyclophilin involved in the process of cell death after organ ischemia-reperfusion. However, the role of CypD in atherosclerosis, especially in necrotic core formation, is unknown.

**Methods:** To clarify the specific role of CypD, apolipoprotein-E/CypD-double knockout (ApoE^-/-^CypD^-/-^) mice were generated. These mice were fed a high-fat diet containing 0.15% cholesterol for 24 weeks to accelerate atherosclerotic lesion development.

**Results:** The deletion of CypD decreased the necrotic core size, accompanied by a reduction of macrophage apoptosis compared to control ApoE^-/-^ mice. In RAW264.7 cells treated with endoplasmic reticulum stress inducer thapsigargin, the release of cytochrome c to the cytosol was attenuated by siRNA-mediated knockdown of CypD. Ly-6C^high^ inflammatory monocytes in the peripheral blood leukocytes and mRNA expression of *Il1b* in the aorta were decreased by the deletion of CypD. In contrast, siRNA-mediated knockdown of CypD did not significantly decrease *Il1b* nor *Ccl2* mRNA expression in RAW264.7 cells treated with LPS and IFN-γ, suggesting that inhibition of inflammation *in vivo* is likely due to decreased cell death in the atherosclerotic lesions rather than a direct action of CypD deletion on the macrophage.

**Conclusions:** This is the first report showing that CypD induces macrophage death and promotes necrotic core formation in advanced atherosclerotic lesions. CypD could be a novel therapeutic target for treating atherosclerotic vascular diseases.

## Introduction

Atherosclerotic cardiovascular diseases, including myocardial infarction and cerebral infarction, are serious clinical problems that threaten patients’ life and cause impairment of quality of life.^1–3^ In advanced atherosclerotic lesions, vulnerable plaques prone to rupture cause these acute thrombotic complications. Thin fibrous caps with decreased vascular smooth muscle cells (VSMCs), lipid-rich macrophage accumulation, and necrotic core formation are characteristic features of vulnerable plaques.^4^ In the necrotic core of advanced atherosclerotic lesions, macrophages, and VSMCs are major cellular components undergoing cell death.^5^ Accumulating evidence from experimental animal studies suggests that impairment of the removal machinery of apoptotic cells in the plaque, i.e., defective efferocytosis, also leads to the accumulation of dead cells and the formation of necrotic cores.^6^ An autopsy study of patients after sudden coronary death shows that necrotic core size was associated with hyperlipidemia and the diabetic state of the patients.^7^ In patients with these two risk factors, macrophage accumulation and necrotic core size were increased. Despite the clinical and experimental evidence showing that necrotic core formation is associated with cardiovascular events, detailed molecular mechanisms by which plaque macrophages result in cell death are largely unknown.

Cyclophilin D (CypD), a mitochondria-specific cyclophilin, contributes to mitochondrial homeostasis by regulating the opening of the mitochondrial permeability transition pore (mPTP). During myocardial ischemia/reperfusion injury, the opening of mPTP and release of cytochrome c to the cytosol induce subsequent cell death.^8, 9^ We have previously demonstrated that cyclophilin inhibitor, cyclosporine A (CsA), attenuates ischemia/reperfusion injury and decrease infarct size in the heart and brain.^10, 11^ It is, however, controversial whether CypD regulates necrotic core formation in advanced atherosclerotic lesions. In animal experiments, non-specific cyclophilin inhibitor CsA has been reported to be atherogenic,^12^ while others indicated that blockade of cyclophilin A function by gene manipulation in mice is atheroprotective.^13^ Therefore, we have planned this study to elucidate the specific role of cyclophilin D in the pathogenesis of atherosclerosis, especially in the necrotic core formation in advanced atherosclerotic lesions. To address this point, we have created genetically altered mice lacking CypD and apolipoprotein-E (ApoE).

## Methods

Please see the Major Resource Table in the Supplemental Materials for additional information about material and methods.

### Experimental animals

The committee on ethics of animal experiments, Kyushu University Graduate School of Medical Sciences, reviewed and approved all experimental animal protocols. Cyclophilin D knockout (CypD^□/-^) mice (Stock No.009071) and ApoE knockout (ApoE^-/-^) mice (Stock No.002052), both of which are C57Bl/6J background, were purchased from The Jackson Laboratory (Bar Harbor, ME). CypD^-/-^ and ApoE^-/-^ mice were crossed to generate ApoE^-/-^CypD^-/-^ double knockout mice. These mice were maintained in the animal facility of Kyushu University under a 12/12-hour light and dark cycle. Mice were fed a γ-irradiated normal laboratory diet (No. FR-1, Funabashi Farm) and water ad libitum. From 8 weeks of age, male ApoE^-/-^ and ApoE^-/-^CypD^-/-^ mice were fed a high-fat diet (HFD) containing 21% fat from lard supplemented with 0.15% cholesterol (Oriental Yeast, Tokyo, Japan) for the indicated period in the Result. Then, mice were euthanized by intravenous administration of 100 mg/kg pentobarbital, and blood samples were collected from the inferior vena cava. After saline perfusion with physiological pressure from the apex of the left ventricle, the aorta, spleen, and fat tissues (epididymal and leg subcutaneous) were dissected for further analysis. Heart rate and blood pressure were measured before sacrifice by a tail-cuff system MK-2000ST (Muromachi Kikai Co., Ltd., Tokyo, Japan). Serum cholesterol concentrations were measured by LabAssay ™ Cholesterol (FUJIFILM Wako Pure Chemical Corporation, Osaka, Japan) according to the manufacturer’s instructions. The number of blood cells, platelets, and hemoglobin concentration were measured using MEK-6450 (NIHON KOHDEN, Tokyo, Japan).

### Histopathology and immunohistochemistry

Excised tissues were fixed with 10% neutral buffered formalin overnight and replaced with 15% and 30% sucrose sequentially. Then, tissues were frozen in the OCT compound (Sakura Finetek Japan Co., Ltd., Tokyo, Japan) and cut by microtome with 7 μm thickness. The aortas at the aortic valve level were stained using an Elastica van Gieson staining kit (Sigma-Aldrich, Inc., St. Louis, MO) and Picrosirius red staining kit (Polysciences, Inc., Warrington, PA) according to the manufacturer’s instruction. In the sections stained with Elastica van Gieson, the plaque area was quantified as an intimal area surrounded by internal elastic lamina except for the lumen area of the aorta. The necrotic core area was determined as previously reported.^14^ Briefly, the area with no extracellular matrix (collagen and elastin fiber) replaced by dead cells/cellular debris was quantified as a necrotic core area. After Picrosirius red staining, sections were observed using polarized microscopy to quantify intraplaque collagen content. All the histopathological analyses were performed by an independent investigator who was blinded to the group of animals. Immunostaining was performed with the following antibodies: mouse anti-cyclophilin F (G-9) (Cat No. sc-376061, Santa Cruz Biotechnology, Inc., Dallas, TX), purified rat anti-Mac-3 (Cat No. 550292, BD Pharmingen, San Diego, CA), rabbit polyclonal anti-α-smooth muscle actin (SMA) antibody (Thermo Fisher Scientific Inc., Waltham, MA), and rabbit monoclonal anti-Ki-67 (Abcam, Cambridge, UK). Staining with mouse-origin primary antibody was performed with a Mouse-on-Mouse detection kit (Vector Laboratories, Inc., Newark, CA) according to the manufacturer’s instruction. Double staining was performed using two substrates, AEC and Vector Blue (Vector Laboratories, Inc.). Apoptotic cells were detected by TUNEL (Terminal deoxynucleotidyl transferase dUTP nick end labeling) staining using *in situ* apoptosis detection kit (Takata Bio Inc., Shiga, Japan) or immunostaining of cleaved caspase-3 (Cell Signaling Technology, Danvers, MA).

### Flow cytometry analysis

Flow cytometric data were acquired using a Gallios cytometer (Beckman Coulter, Fullerton, CA. USA). For peripheral blood samples, red blood cells were removed by hemolysis (VersaLyse, Beckman Coulter) and centrifugation at 240 × g for 5 minutes at 4°C. The remaining leukocytes were washed with Hanks’ balanced salt solution (HBSS). Cells were incubated with fluorochrome-labeled monoclonal antibodies after blocking the Fc receptor with anti-CD16/32 monoclonal antibodies (BD Pharmingen, San Diego, CA. USA). Data were analyzed with FlowJo or Kaluza software. Monocytes/macrophages were identified as CD11b^high^ lineage markers (includes Ly6G, NK1.1, CD90, CD49b, B220)^low^ Ly6C^high/low^. The following antibodies were used: anti-CD90, 53-2.1 (BD Pharmingen), anti-B220, RA3-6B2 (BD Pharmingen), anti-CD49b, DX5 (BD Pharmingen), anti-NK1.1, PK136 (BD Pharmingen), anti-Ly6G, 1A8 (BD Pharmingen), anti-CD11b, M1/70 (BD Pharmingen), anti-Ly6C-FITC, AL-21 (BD Pharmingen), anti-mouse CD147, OX-114 (BioLegend, San Diego, CA. USA), anti-mouse CD206, C068C2 (BioLegend).

### Semi-quantitative real-time PCR

RNA was extracted from tissues or cells by illustra RNAspin Mini kit (G.E. Healthcare, PA. USA), and cDNA was synthesized using PrimeScript RT reagent Kit (Perfect Real Time, TAKARA BIO Inc. Shiga, Japan). The mouse aorta was homogenized by TissueLyser II (Qiagen, Venlo, Netherlands) according to the manufacturer’s instructions, and then mRNAs were extracted. Semi-quantitative real-time PCR was performed with SYBR Premix DimerEraser (Perfect Real Time, TAKARA BIO Inc.) and a StepOnePlus real-time PCR system (Applied Biosystems, MA. USA). Primer designs are listed in Table 1. Data were calculated by ΔΔCT method and expressed in arbitrary units normalized by β-actin.

**Table 1.**
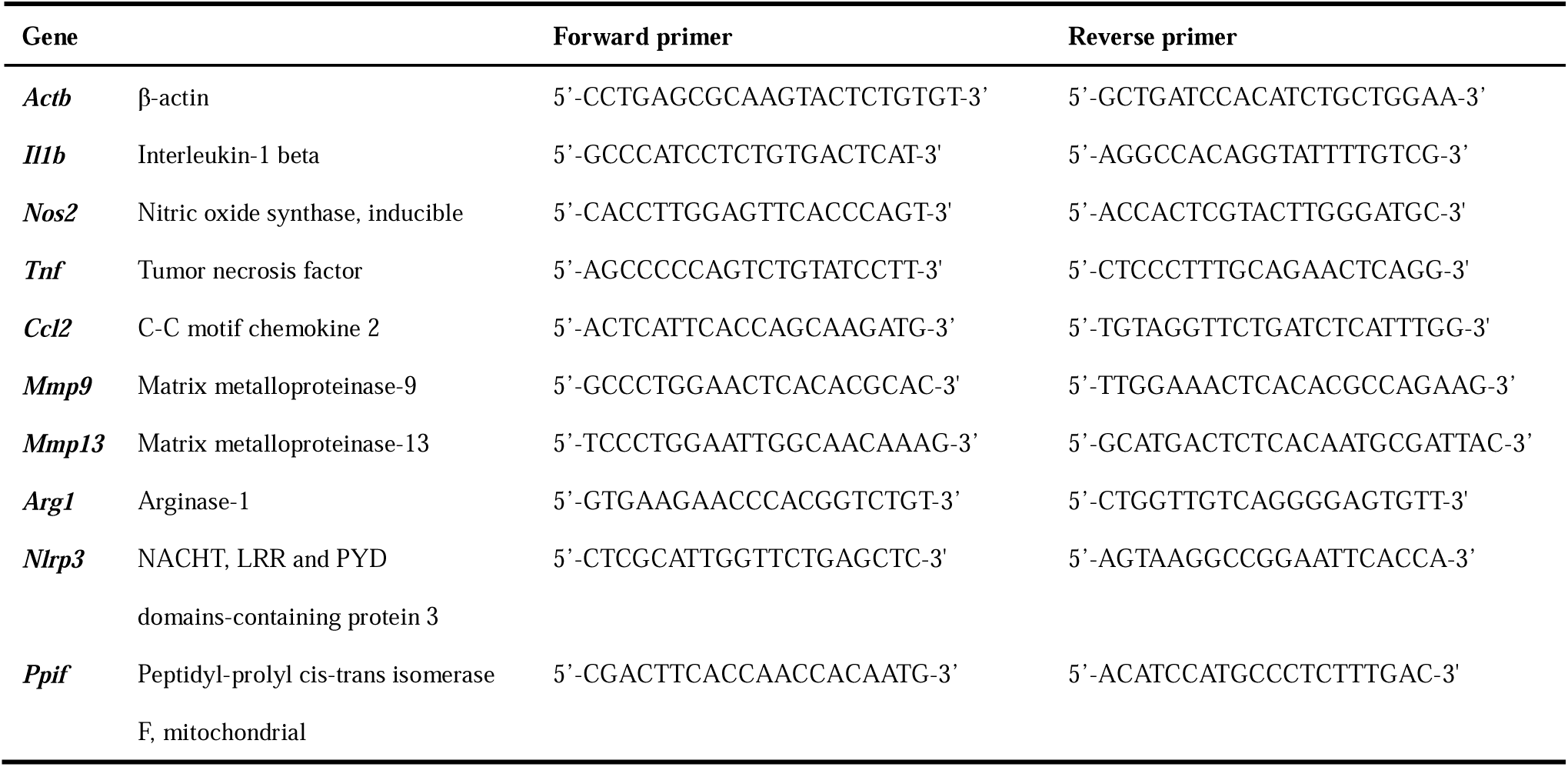
Primer design for semi-quantitative RT-PCR.

### *Ex vivo* macrophage activation

Peritoneal macrophages elicited by Brewer modified thioglycolate medium (Beckton Dickenson, Franklin Lakes, NJ) in donor mice, and RAW264.7 cells (TIB-71, ATCC, Manassas, VA) were used for *ex vivo* or *in vitro* culture experiments. These cells were cultured with RPMI 1640 medium (R8758, Lonza, Basel, Switzerland) containing 10 % fetal bovine serum (FBS) (12003C, Sigma-Aldrich, St. Louis, MO) and 1% penicillin/streptomycin. Classically activated macrophages were prepared by stimulating macrophages with 10 ng/mL lipopolysaccharide (LPS) (L5293, SigmalAldrich) and 10 ng/mL mouse interferon-γ (mIFN-γ, 11276905001, Roche, Basel, Switzerland). In loss-of-function experiments, 30 nM of CypD (*Ppif*) siRNA or an equivalent concentration of control non-targeting siRNA (Horizon Discovery Ltd., Cambridge, UK) was transiently transfected with SilenceMag (SM10200, OZ Biosciences, Marseille, France) according to the manufacturer’s instruction. In gain-of-function experiments, the expressing vector of CypD or control vector (GeneCopoeia, Rockville, MD) was transiently transfected into RAW264.7 cells using PolyMag Neo (PG60200, O.Z. Biosciences). Culture supernatant was exchanged 24 hours later and used for experiments.

### Quantification of cytochrome c in the cytosolic fraction

RAW264.7 cells transfected with 30 nM CypD (*Ppif*) siRNA or non-targeting control siRNA in the same manner as above were pretreated with 1.0 μM thapsigargin for 24 hours. According to the manufacturer’s instructions, cytosolic fractions were isolated to measure cytochrome c concentration using a mitochondrial isolation kit for culture cells (ThermoScientific). The cytochrome c concentrations in the isolated cytosolic fraction were measured by rat/mouse cytochrome c quantikine ELISA Kit (MCTC0, R&D Systems, Inc., Minneapolis, MN).

### Statistics

The GraphPad Prism software program, version 9 (GraphPad Software, La Jolla, CA), was used for statistical analyses. All data were reported as the mean±SEM. The data were first analyzed for normality using the Shapiro-Wilk test and equal variance using F-test. Data that did not satisfy these tests were analyzed by the Mann-Whitney test. Other statistical tests comparing the two groups were performed by Student’s t-test. Statistical analyses of the *in vitro* loss-of-function and gain-of-function experiments were performed by a 2-way ANOVA with Bonferroni’s post-test. Probability values <0.05 were considered statistically significant.

## Results

### CypD deficiency increased serum cholesterol levels after 24 weeks of a high-fat diet

ApoE^-/-^ mice and ApoE^-/-^CypD^-/-^ mice were fed HFD containing 0.15% cholesterol for 24 weeks to accelerate atherosclerotic lesion development. In these mice, body and spleen weights were heavier in ApoE^-/-^CypD^-/-^ mice, accompanied by increased serum cholesterol levels compared to ApoE^-/-^ mice (Table 2). Therefore, we have quantified body weight and food intake until 12 weeks after HFD feeding. Until this point, deletion of CypD showed no statistically significant effects on body weight, food intake (Figures 1A and 1B), and serum cholesterol levels (1378±137 vs. 1675±102 mg/dL) as compared to control ApoE^-/-^ mice. There were no significant differences in the number of white blood cells, platelets, and hemoglobin concentration in the peripheral blood, too (Table 2). Hemodynamic parameters, including heart rate and systolic blood pressure, were comparable between these 2 groups. There were no significant differences in the weight of other organs and tissues, e.g., liver, heart, epididymal visceral fat tissue, and leg subcutaneous fat tissue. Immunohistochemical analysis demonstrated the expression of CypD in plaque macrophages, especially foam cells in subendothelial space and macrophages surrounding the necrotic core (Figure 1C).

**Figure 1.**
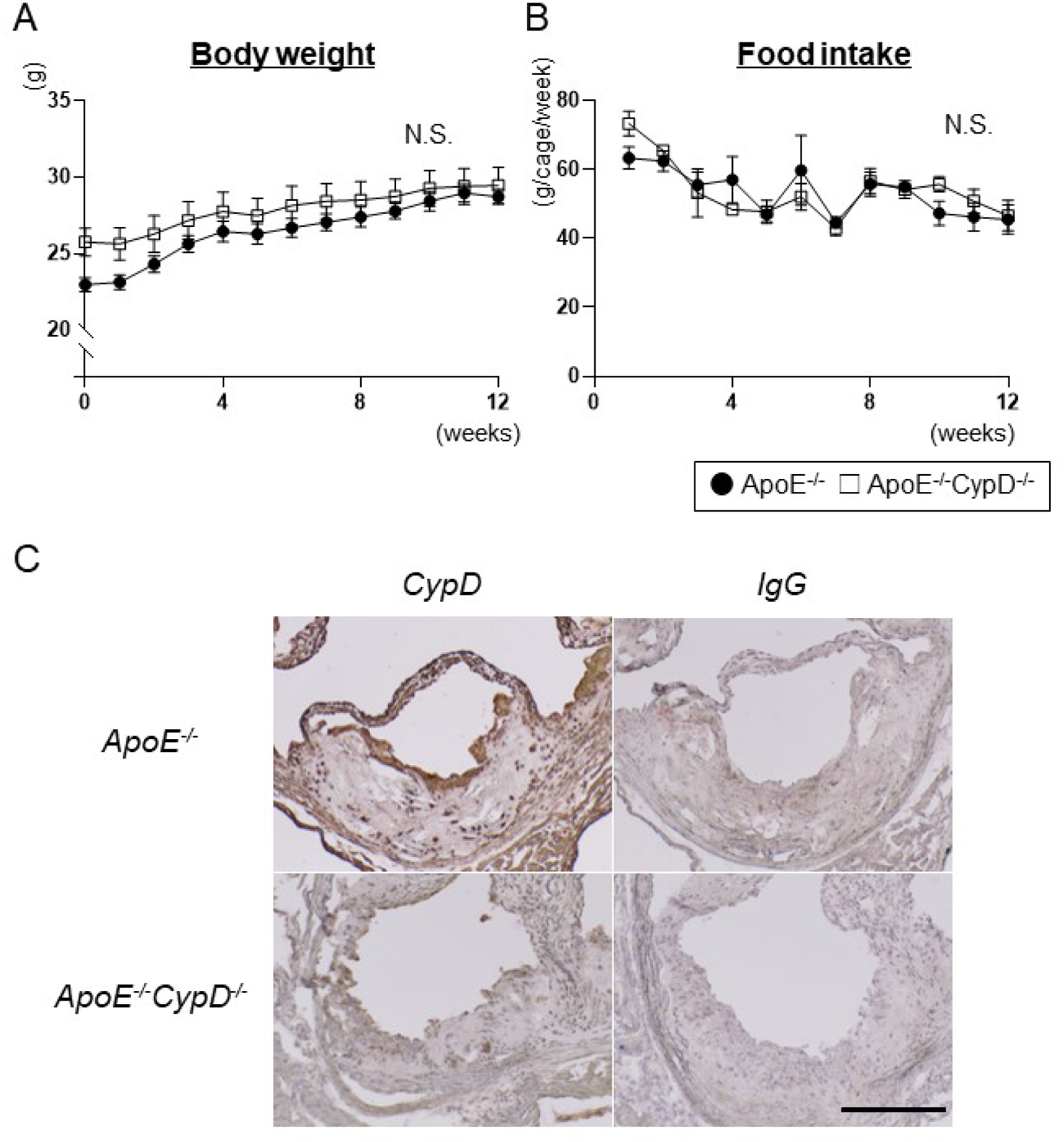
Mouse model of atherosclerosis in cyclophilin D(CypD)-deficient mice. A and B, the time course of body weight and food intake of ApoE^-/-^ and ApoE^-/-^CypD^-/-^ mice. X-axes indicate weeks after the beginning of a high-fat diet. N=8 each. Y-axes indicate body weight (A) and food intake (g/cage/week). Four mice were kept in each cage, and food intake was measured as the total intake of the four mice; Error bars indicate SEM between cages. C, immunostaining with anti-CypD antibody revealed expression of CypD in plaque macrophages of ApoE^-/-^ mice after 24 weeks of a high-fat diet. CypD; anti-CypD antibody, IgG; non-immune control IgG, ApoE^-/-^; Apolipoorotein-E knockout mouse, ApoE^-/-^CypD^-/-^; ApoE/CypD double knockout mice, NS; not significant. The scale bar indicates 200 μm.

**Table 2.** Body weight, organ weight, hemodynamic parameters, and serum lipid profile of ApoE^-/-^ and ApoE^-/-^CypD^-/-^ mice.

|  | *ApoE<sup>-/-</sup>* | *ApoE<sup>-/-</sup>CypD<sup>-/-</sup>* | p-value |
| --- | --- | --- | --- |
| Body weight (g) | 34.73±1.37 | 40.26±1.34 | <0.01 |
| Liver weight (mg) | 1670±169.1 | 1812±119.6 | ns |
| Spleen weight (mg) | 124.2±10.96 | 164.3±13.5 | <0.05 |
| Visceral fat weight (mg) | 205.1±8.91 | 219.5±7.25 | ns |
| Subcutaneous fat weight (mg) | 378.7±81.18 | 489.5±65.47 | ns |
| Heart rate (/min) | 682±22.81 | 770.0±34.95 | ns |
| Systolic blood pressure (mmHg) | 113.3±7.96 | 102.3±5.14 | ns |
| Hemoglobin (g/dl) | 11.23±0.19 | 11.99±0.35 | ns |
| White blood cells (x10<sup>2</sup>/μl) | 88.2±12.3 | 73.8±12.7 | ns |
| Platelets (x10<sup>4</sup>/μl) | 86.32±14.08 | 111.7±8.5 | ns |
| Total cholesterol (mg/dl) | 936.3±82.19 | 1868±153.1 | <0.0001 |

3 All the parameters were measured after 24 weeks of high-fat diet feeding. N=3-16.

### CypD promotes necrotic core formation in advanced atherosclerotic lesions

ApoE^-/-^ and ApoE^-/-^CypD^-/-^ mice were fed an HFD to accelerate atherosclerotic lesion development. After 24 weeks of HFD feeding, pathological features associated with advanced atherosclerotic lesions, including necrotic core formation and accumulation of VSMCs and macrophages, were observed. The necrotic core area in aortic root plaques, determined as acellular and anuclear plaque area containing cholesterol clefts, was significantly decreased by the deletion of CypD (Figure 2). The atherosclerotic plaque area was also significantly decreased by the deletion of CypD. Collagen content quantified by picrosirius red staining observed under polarized microscopy was comparable between the 2 groups. In immunohistochemical analyses, macrophage content determined as a ratio of Mac-3-positive area relative to the plaque area was increased in ApoE^-/-^CypD^-/-^ mice. However, the absolute Mac-3-positive area was not significantly affected (ApoE^-/-^; 3.46±0.37 vs ApoE^-/-^CypD^-/-^; 2.63±0.35 ×10^5^ μm^2^, N=8, NS). VSMCs determined as α-SMA positive cells in the plaque were significantly decreased.

**Figure 2.**
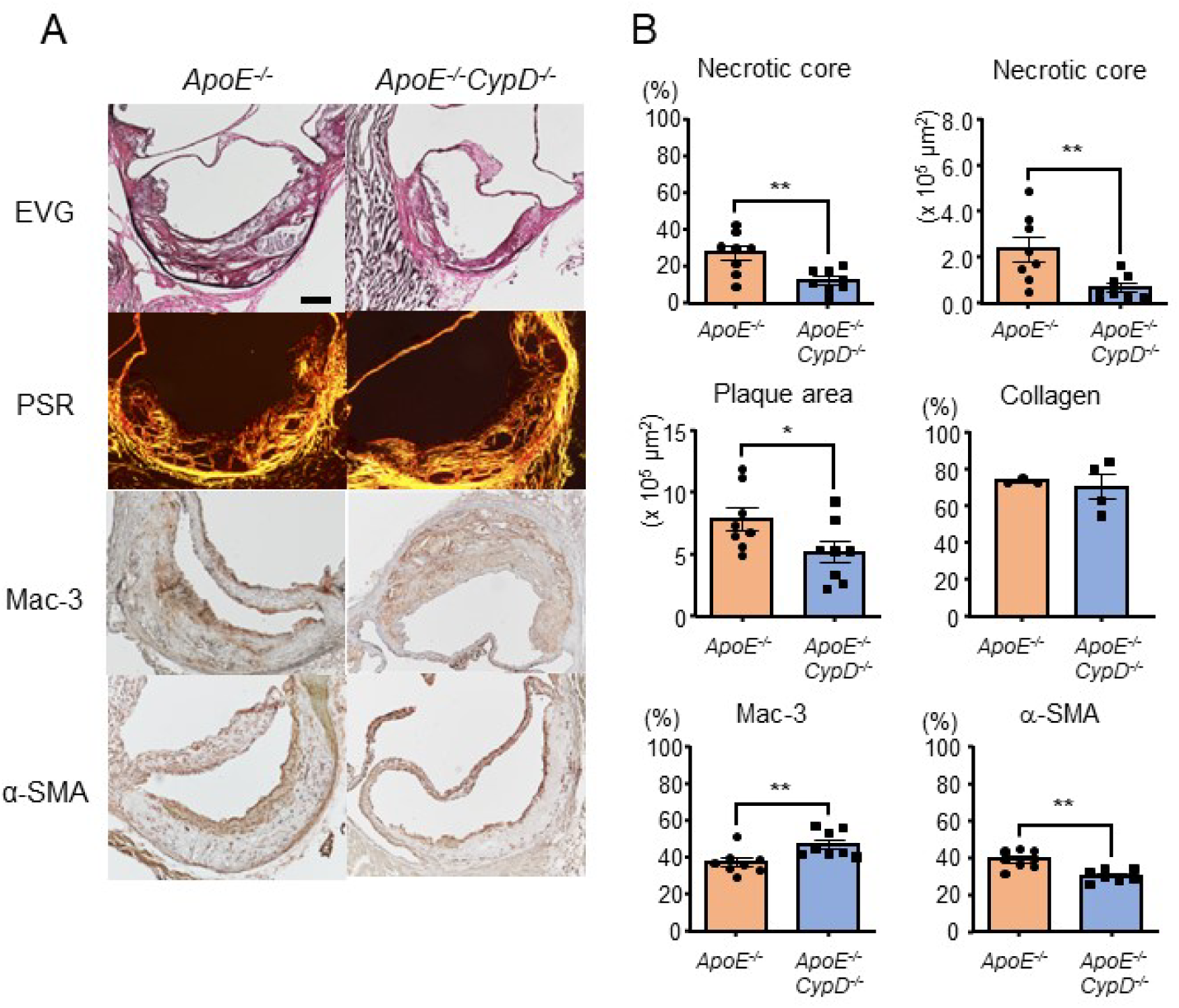
Histopathological analyses of the aortic lesion after 24 weeks of high-fat diet feeding. A, Cross sections of the aortic root. The necrotic core area was determined as the acellular area in aortic valve sections stained by EVG. EVG; Elastica van Gieson staining, PSR: Picrosirius Red staining. The scale bar indicates 100 μm. B, Quantitative data of necrotic core and plaque area were quantified in sections stained with EVG. Collagen areas were quantified in sections stained by PSR. Mac-3 and α-SMA were measured as brown areas stained by immunostaining of Mac-3 and α-SMA, respectively, relative to the total plaque area. N=3-8. *p<0.05, **p<0.01.

### Blockade of CypD inhibits apoptotic cell death in macrophages

To examine the role of CypD in cell proliferation and death, Ki-67, TUNEL+ cells, and cleaved caspase-3 were stained in the aorta sections at the aortic valve level. Ki-67 staining demonstrated proliferating cells in the intima of atherosclerotic lesions (Figure 3A left, arrowheads). The proliferating cells were significantly decreased in ApoE^-/-^CypD^-/-^ mice compared to ApoE^-/-^ mice. TUNEL^+^ apoptotic cells were also observed in the intima of plaques, especially around the necrotic core lesions (Figure 3A right, arrowheads). In these lesions, the deletion of CypD significantly decreased TUNEL^+^ cells. To further examine whether macrophage apoptosis is decreased by CypD deletion, double immunostaining of cleaved caspase-3 and Mac-3 was performed. As shown in Figure 3B, the deletion of CypD significantly decreased the ratio of cleaved caspase-3 stained cells (red-brown) in total macrophages (blue). In contrast, efferocytosis, determined as the ratio of cleaved caspase-3 stained cells colocalized with Mac-3 positive cells relative to cleaved caspase-3 stained cells not colocalized with Mac-3 stained cells,^15^ was not significantly decreased by deletion of CypD.

**Figure 3.**
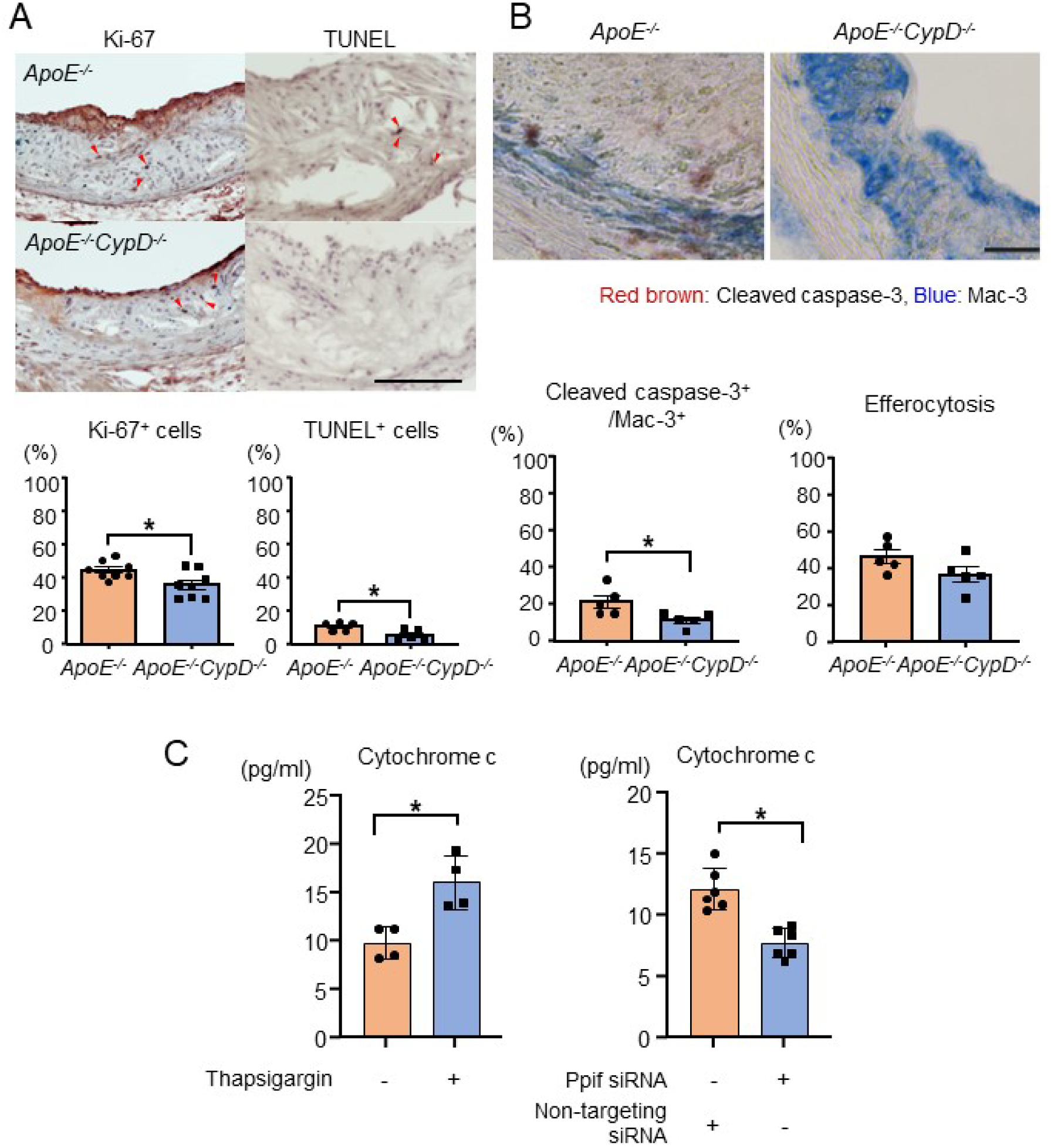
Histopathological analyses of proliferating and apoptotic cells in advanced atherosclerotic lesions. A, proliferating, and apoptotic cells were stained by anti-Ki-67 antibodies and TUNEL. The graphs show quantitative data demonstrated as percent positive cells in the intima of atherosclerotic plaques. N=5-8. The scale bar indicates 100 μm. B, Double immunostaining of cleaved caspase-3 (red-brown) and Mac-3 (blue) to evaluate the apoptotic cells and efferocytosis. Efferotycosis was quantified as a ratio of apoptotic cells that colocalized with Mac-3 positive cells relative to total apoptotic cells. N=5, each. The scale bar indicates 100 μm. *p<0.05. C, ELISA of cytosolic cytochrome c in RAW264.7 cells treated with thapsigargin for 24 hours. N=6. *p<0.05.

To elucidate whether CypD regulates apoptotic cell death in macrophages, we have performed an *in vitro* experiment using a cultured macrophage cell line, RAW264.7. In RAW264.7 cells, endoplasmic reticulum (ER) stress-induced apoptotic cell death was induced by thapsigargin, an inhibitor of Ca^2+^-ATPase on the ER membrane. In this condition, cytochrome c concentration in the cytosol fraction was measured. Leakage of cytochrome c from the mitochondria to the cytosol via mPTP opening activates the downstream apoptosis pathway, including activation of caspase-9 and caspase-3. As shown in Figure 3C, siRNA-mediated knockdown of CypD (Ppif) suppressed the release of cytochrome c to the cytosol in thapsigargin-treated RAW264.7 cells, suggesting that CypD promotes ER stress-induced apoptotic cell death in macrophages.

### Deletion of CypD decreases Ly-6C^high^ monocytes in the peripheral blood, and *Il1b* expression in the aorta of ApoE^-/-^ mice

Monocyte/macrophage-mediated inflammation plays an essential role in the formation and progression of atherosclerosis.^16, 17^ Therefore, we have performed flow cytometry analysis to evaluate the monocyte skewing to inflammatory phenotype in the peripheral blood. These inflammatory monocytes are the source of macrophages that accumulates in the plaques. There was no significant difference in the cell count of the CD11b^+^Lin^-^ monocyte population between ApoE^-/-^ and ApoE^-/-^CypD^-/-^ mice in the circulating leukocytes, but the percentage of Ly-6C^high^ monocytes in this population was significantly decreased in the peripheral blood of ApoE^-/-^CypD^-/-^ mice (Figure 4).

**Figure 4.**
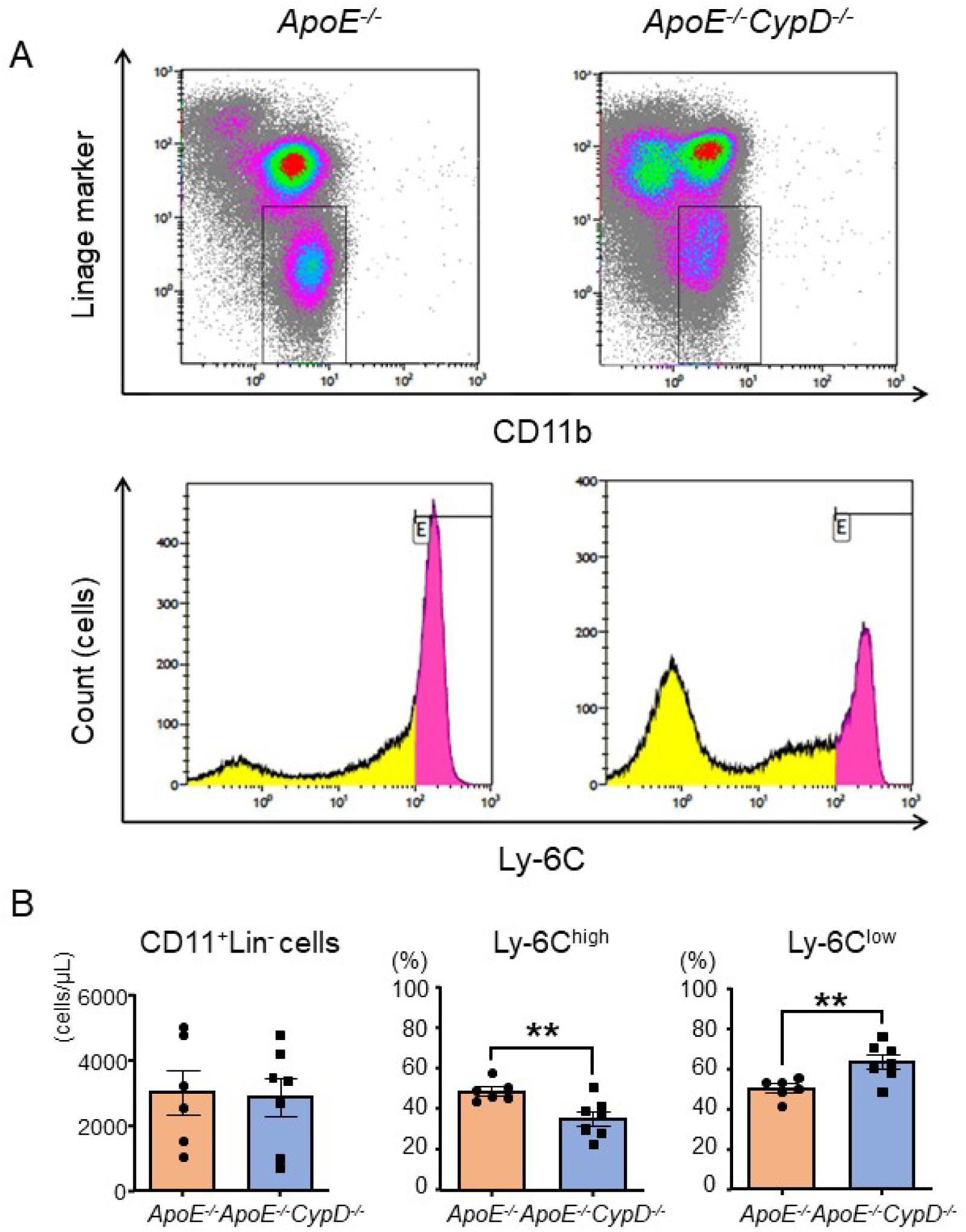
Flow cytometry analysis of circulating monocytes in the peripheral blood. A, peripheral blood cells were collected, and monocytes were gated as CD45^+^Lin^-^CD11b^+^ cells (above 2 panels). The lower 2 panels show the histograms of Ly-6C staining. The ly-6C^high^ population was demonstrated in pink color. B, quantification of total monocyte number, the ratio of Ly-6C^high^ cells, and Ly-6C^low^ cells in circulating monocytes. **p<0.01. N=6 or 7.

To further investigate the effects of CypD deletion on inflammation, we quantified mRNA expression levels in the aortas of ApoE^-/-^ mice fed a HFD for 12 weeks. As shown in Figure 5, mRNA expression of *Il1b* was significantly suppressed by the deletion of CypD. Reduction/induction of other molecules related to macrophage activation or extracellular matrix degradation, including *Mmp9*, *Mmp13*, *Arg1*, and *Nlrp3,* did not reach significant differences.

**Figure 5.**
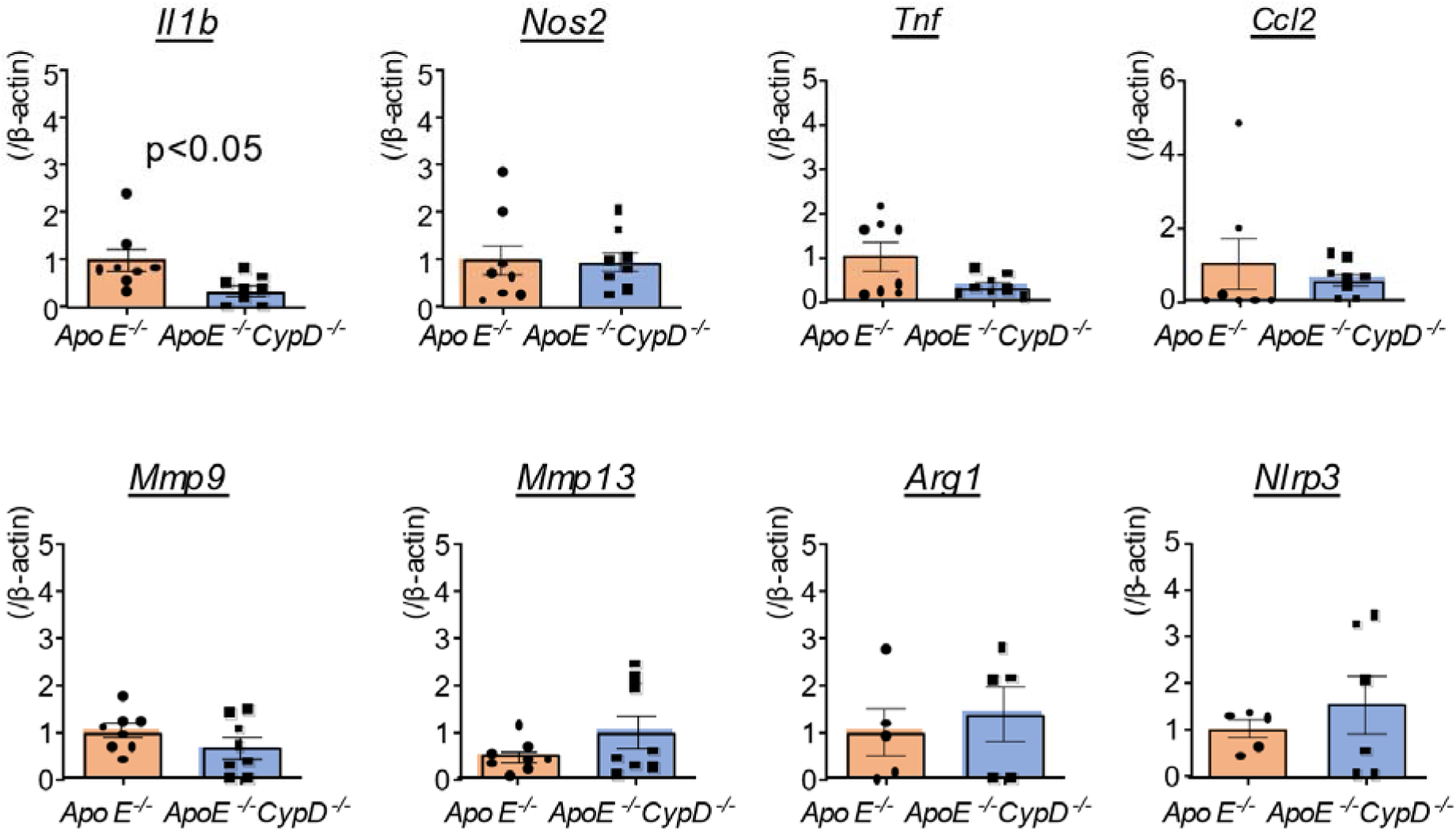
Semi-quantitative PCR of the aorta harvested from ApoE^-/-^ and ApoE^-/-^CypD^-/-^ mice. mRNA was extracted from the aorta of ApoE^-/-^ and ApoE^-/-^CypD^-/-^ mice after 12 weeks of HFD, and mRNA expressions of molecules associated with macrophage activation or extracellular matrix degradation were quantified. Mmp; matrix metalloproteinase, Nrlp3; NLR family pyrin domain containing 3. N=5-8.

### CypD did not induce the expression of molecules associated with inflammatory macrophages

The result of flow cytometry and semi-quantitative real-time PCR analyses raised a possibility that CypD promotes macrophage activation. Hence, we have performed *in vitro* loss-of-function and gain-of-function studies to elucidate the direct and specific role of CypD in macrophage polarization. In the murine macrophage cell line, RAW264.7, siRNA-mediated knockdown of CypD (*Ppif*) did not significantly decrease or increase the expression of inflammatory cytokines, including *Il1b* and *Ccl2* (Figure 6A). mRNA expressions of *Tnf*, *Nos2*, and *Il-6* were not significantly affected. In contrast, transient overexpression of *Ppif* using a plasmid vector decreased *Il-1*β mRNA, but the expression of *Ccl2* was not significantly modified. Expression of *Arg1* was not significantly modified by transient overexpression of *Ppif*, either (Figure 6B). These results suggest that CypD does not directly regulate the differentiation of macrophages into inflammatory or alternatively activated macrophages.

**Figure 6.**
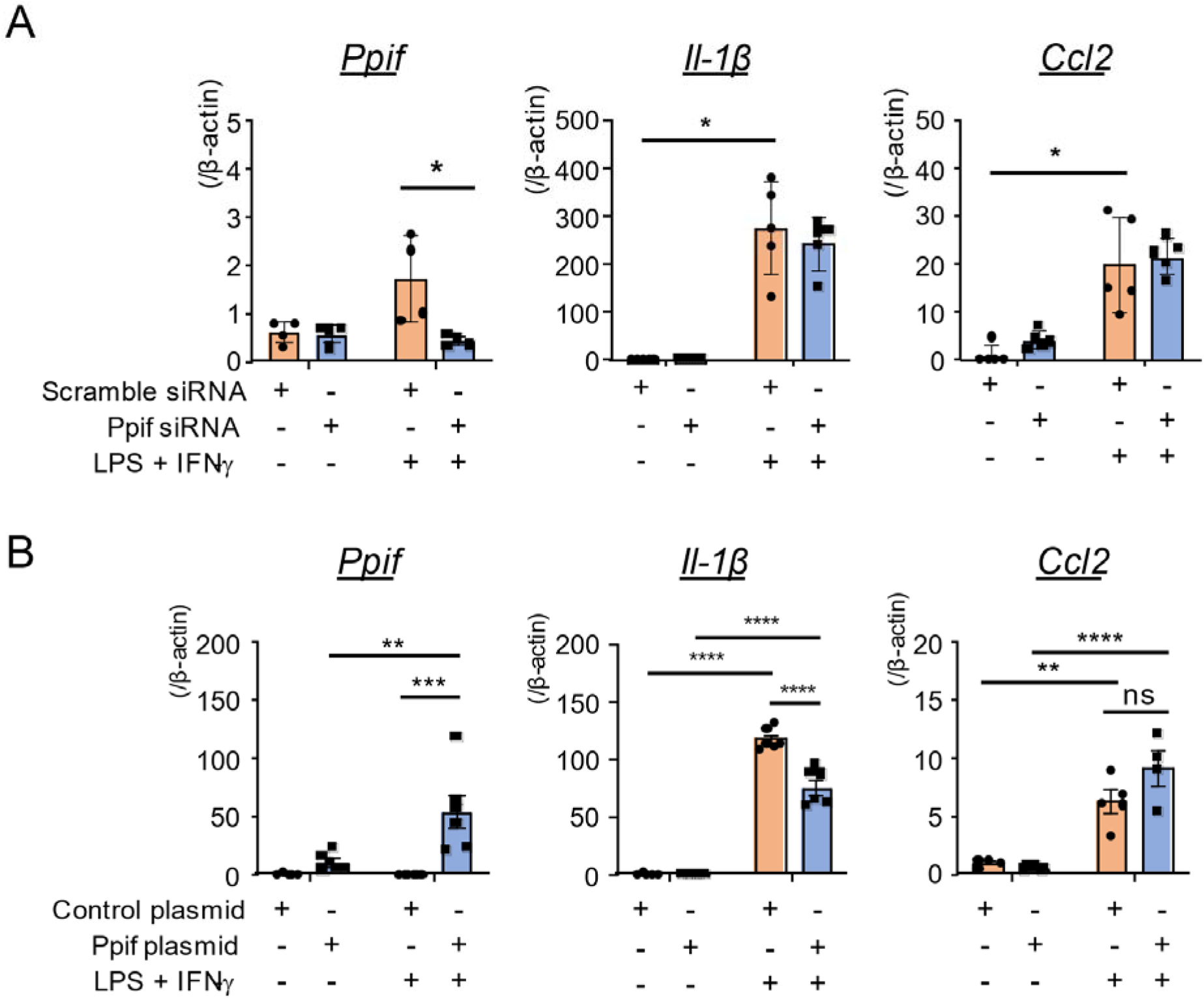
Loss-of-function and gain-of-function experiments in RAW264.7 cells. A, CypD (*Ppif*) mRNA was knocked down by siRNA transfected with Magnetofection. These cells were stimulated with LPS and IFN-γ 48 hours later, and then mRNA was extracted for quantitative analysis. N=5. *p<0.05. B, the expression vector of Ppif was transiently transfected to RAW264.7 cells, and mRNA expression was quantified 48 hours later. **p<0.01, ***p<0.001, ****p<0.0001. N=5-6.

## Discussion

Here, we have demonstrated that deletion of CypD inhibits necrotic core formation in a mouse model of atherosclerosis for the first time. Immunohistochemistry indicated the expression of CypD in plaque macrophages. In these mice, deletion of CypD decreased necrotic core size, accompanied by increased macrophage accumulation. Immunostaining of Ki-67, however, indicates the reduction of cell proliferation in the plaque of ApoE^-/-^CypD^-/-^ mice. This discrepancy, i.e., macrophage accumulation was increased despite the reduction of their proliferation, suggested that reduction of cell deaths, macrophages are a major source of these cells,^18^ was a major mechanism of increased macrophage accumulation in ApoE^-/-^CypD^-/-^ mice. The result of TUNEL staining showing decreased apoptosis in the plaque underpins the above speculation. Flow cytometry of the peripheral blood indicated the reduction of circulating inflammatory, i.e., Ly-6C^high^, monocytes in ApoE^-/-^CypD^-/-^ mice. Semi-quantitative real-time PCR of the aorta indicated the reduction of *Il-1*β mRNA expression in the aorta of ApoE^-/-^CypD^-/-^ mice. These results suggest that the deletion of CypD attenuates inflammation in ApoE^-/-^ mice. However, *in vitro* loss-of-function and gain-of-function experiments using mouse macrophages suggested that CypD does not directly regulate macrophage polarization toward inflammatory subsets. Therefore, the direct role of macrophage CypD in macrophage polarization is speculated to be scant.

Plaque rupture of vulnerable plaque is a major cause of acute thrombotic complications in coronary arteries. Clinical studies using intravascular ultrasound demonstrated that necrotic core is often observed in lipid-rich plaques, which are prone to rupture compared to fibrous atheroma.^19, 20^ CsA, a clinically available agent used in patients after heart transplantation, forms a complex with cyclophilin and inhibits calcineurin activity. However, there is insufficient clinical evidence of whether inhibition of cyclophilins decreases necrotic core formation and subsequent acute coronary events because its use is limited to patients after organ transplantation due to its potent immunosuppressive action and undesired side effects, including renal dysfunction, hypertension, and dyslipidemia. Our data indicates a possibility that pharmacological inhibition of CypD decreases necrotic core formation and plaque rupture in patients, but local drug delivery using a nanotechnology-based system or drug-eluting stents may be necessary to avoid systemic side effects. We have previously demonstrated that nanoparticle-mediated delivery of cyclosporin inhibits ischemia-reperfusion injury in the heart and brain.^10, 11^ These modalities enable efficient drug delivery to plaque macrophages. Hence, it is desired that further technological innovation will make it possible to apply these technologies to prevent cardiovascular events in chronic atherosclerosis patients.

Detailed molecular mechanisms about how CypD regulates cell death in atherosclerotic plaque are largely unknown. In reperfusion injury after organ ischemia, CypD regulates the opening of mPTP and causes a release of cytochrome c to the cytosol, which induces subsequent cell death.^8, 9^ In the necrotic core, it has been reported that apoptosis is one of the major mechanisms of cell death.^18^ Our data showing the reduction of TUNEL^+^ or cleaved caspase-3^+^ cells in advanced atherosclerotic lesions of ApoE^-/-^CypD^-/-^ mice indicates apoptosis is, at least in part, one of the cell death mechanisms in the necrotic core. It is, however, unknown whether similar mechanisms, i.e., mPTP opening, promote the process of macrophage death in advanced atherosclerotic lesions. Instead, experimental data indicates ER stress-mediated mechanisms are important in the cell death process in atherosclerotic lesions.^21^ Especially, free cholesterol-induced changes of the ER membrane, impairment of ER function as a calcium reservoir, and activation of unfolded protein responses are important molecular mechanisms underlying the cell death process.^22^ Clinical data also indicate an association between ER stress, apoptosis, and plaque vulnerability. Miyoshi M et al. reported that transcriptional factor CHOP, which is induced by ER stress, is increased in atherosclerotic lesions with thin fibrous caps and abundant TUNEL^+^ cells. In cultured VSMCs and macrophages, they have demonstrated that 7-ketocholesterol, a major oxidation product, induced ER stress and cell death, both of which were attenuated by siRNA-mediated knockdown of CHOP.^23^ Here, we have demonstrated that the knockdown of CypD suppresses the thapsigargin-induced leakage of cytochrome c. In combination with the *in vivo* results, it is suggested that genetic deletion of CypD decreases apoptosis of macrophages in advanced atherosclerotic lesions.

Monocyte/macrophage-mediated inflammation promotes atherosclerosis from the initial plaque formation to plaque rupture in advanced lesions resulting in acute coronary syndromes. In ApoE^-/-^ mice, deletion of CypD decreased plaque size. Therefore, flow cytometry analysis and quantification of inflammatory cytokines were performed to examine the role of CypD in inflammation. Flow cytometry demonstrated a reduction of Ly-6C^high^ inflammatory monocytes, and PCR demonstrated a reduction of *Il-1*β mRNA expression in ApoE^-/-^CypD^-/-^ mice. In contrast, *in vitro* loss-of-function and gain-of-function experiments, which were performed to elucidate the direct role of CypD in macrophages, showed no significant effects on macrophage polarization except the reduction of *Il-1*β mRNA in Ppif plasmid transfected macrophage cell line. These results suggest that inflammation *in vivo* was suppressed in CypD-deficient as a result of decreased monocytes/macrophage cell death, although there are possibilities that CypD in other types of cells, including endothelial cells and VSMCs, play critical roles in plaque inflammation.^24^ In necrotic cores, the retention of dead cells could induce inflammation by releasing several proinflammatory molecules, such as damage-associated molecular patterns (DAMPs).^25^ In this decade, molecular mechanisms of other forms of cell death, e.g., necroptosis, ferroptosis, and pyroptosis, have been clarified. These points raise a possibility that CypD regulates these types of cell death. However, the results of TUNEL staining suggest that the anti-apoptotic effect contributed to the reduction of necrotic core in *in vivo* settings.

There are several steps to overcome for the clinical translation of the results shown here. At first, the clinically available cyclophilin inhibitor, CsA, has a potent immunosuppressive action and is hardly used to treat chronic lifestyle-related diseases. Furthermore, long-term use of CsA could cause several undesired effects, including renal dysfunction and lipid metabolism disorders. In ApoE^-/-^CypD^-/-^ mice, long-term HFD loading caused increased serum cholesterol levels and body weight. In contrast, food intake was similar until 12 weeks after the beginning of HFD. The following mechanisms are postulated as mechanisms of cholesterol elevation by CsA: 1) inhibition of cholesterol transport to the intestine by bile acid, 2) binding to circulating LDL-C, and 3) inhibition of lipoprotein lipase activities.^26, 27^ Since free cholesterol has been reported to induce ER stress and apoptosis,^21^ it is assumed that elevated cholesterol per se promoted the necrotic core formation. Nevertheless, the deletion of CypD inhibited necrotic core formation. This result suggests that CypD is a promising therapeutic target for decreasing necrotic core formation.

In conclusion, CypD induces necrotic core formation by promoting mitochondria-mediated macrophage apoptosis in advanced atherosclerotic lesions, which promotes plaque vulnerability. A combination of CsA and drug delivery system or delivery of CypD-targeting nucleic acid might contribute to developing novel and clinically feasible therapeutics that prevent acute thrombotic complications of atherosclerosis and improve patients’ lives.

## Acknowledgments

We thank Tomoko Fujita and Kozue Nakamura for their excellent technical assistance.

## Highlights

- The necrotic core formation is one of the features of vulnerable plaques. oxLDL-induced ER stress is one of the upstream mechanisms, but detailed molecular mechanisms by which cell death occurs in advanced atherosclerotic lesions remain to be elucidated.
- In ApoE^-/-^ mice fed a high-fat diet, deletion of CypD, a molecule regulating mPTP opening, deceased the necrotic core size accompanied by a reduction of macrophage apoptosis.
- In macrophage cell line RAW264.7, ER stress inducer thapsigargin induced cytosolic releases of cytochrome c, which was attenuated by siRNA-mediated knockdown of CypD.
- In RAW264.7 cells, CypD loss-of-function or gain-of-function did not show significant effects on macrophage skewing to the inflammatory phenotype, suggesting that CypD induces cell death independently with polarity shift of macrophages.

